# Translation-driven temporal control for intertwined protein assembly

**DOI:** 10.1101/2025.08.25.672138

**Authors:** Katharina Till, Vanda Sunderlikova, Frank Tippmann, Predrag Jevtić, Matilde Bertolini, Kai Fenzl, Jaro Schmitt, Alexandros Katranidis, Bernd Bukau, Günter Kramer, Michael Rapé, Sander Tans

**Affiliations:** AMOLF, Autonomous Matter Department, Science Park 104, 1098 XG Amsterdam, The Netherlands; Center for Molecular Biology of Heidelberg University (ZMBH), Im Neuenheimer Feld 329, Heidelberg 69120, Germany; Department of Molecular and Cell Biology, University of California at Berkeley, Weill Hall, #3200, Berkeley, CA 9472, USA; Howard Hughes Medical Institute, University of California at Berkeley, Berkeley, CA 9472, USA; Addition Therapeutics, 201 Haskins way, South San Francisco, 94080 California, USA; European Molecular Biology Laboratory, Genome Biology Unit, Meyerhofstrasse 1, Heidelberg, Germany, 69117; Ernst-Ruska Centre for Microscopy and Spectroscopy with Electrons/ER-C-3 Structural Biology, Forschungszentrum Jülich (FZJ), Jülich, Germany; California Institute for Quantitative Biosciences (QB3), University of California at Berkeley, 174 Stanley Hall, #3220, Berkeley, CA 9472, USA; Bionanoscience Department of Delft University of Technology and Kavli Institute of Nanoscience Delft, 2629 HZ Delft, The Netherland

## Abstract

Protein complexes are essential to cells. However, how structurally intertwined protein subunits can assemble faithfully is poorly understood. Here, we reveal a “temporal control” mechanism driven by coupled ribosomes to form intertwined dimers. Using Disome Selective Profiling and optical tweezers, we show that the BTB domains of KEAP1, KLHL12, and PATZ1 form stable closed states as monomers, thus impeding proposed domain-swapping assembly routes. By contrast, the timed emergence of nascent chain segments during translation enables alternative folding-assembly pathways that bypass the closed monomeric state. Analysis indicates that this mechanism works in concert with dimerization quality control by the E3 ligase SCF-FBXL17, and is relevant across the BTB domain family. This study shows that ribosome cooperation expands the range of possible protein architectures.

## Introduction

Protein complex formation is pivotal to all main cellular functions^1–4^. The assembly of two proteins has long been characterized as a simple binding event^5,6^. However, many complexes within the proteome exhibit highly intricate protein-protein interfaces and even intertwined polypeptide chains^7^. Examples range from large nuclear pore complexes^8^, to E3 ubiquitin ligase Cullin complexes^9^, down to interleukin dimers^10^. These features imply major assembly challenges: protein subunits must either be kept partially unfolded until they interact, or initially adopt compact monomeric structures that require remodeling to assemble^11–13^. Moreover, the involved cytosolic exposure of assembly interfaces poses intrinsic risks, with many proteins subunits prone to aggregation^14–16^. These observations suggest that a vast space of critical folding and assembly mechanisms remains unexplored.

We recently showed that over 800 homodimers assemble co-translationally in human cells^17^. In this process termed co-co assembly, dimerization is driven by interactions between nascent chains emerging from nearby ribosomes translating the same or different RNA messages^18,19^. BTB proteins were identified as a main co-co assembly class (**Fig. 1A**). Human cells contain about 200 different proteins with BTB dimerization domains^20^. They perform functions ranging from gene regulation to actin stabilization, and often contain BTB domains in combination with domains such as Kelch, ion transport, and Zinc finger domains^21^. BTB domains are thought to dimerize by “domain-swapping”^20^. Here, a β-strand that is folded in the monomeric state must dissociate to instead fold onto a dimerization partner, giving rise to an intertwined dimer^20,22^ (**Fig. 1B**). However, the involved energetic cost and aggregation risks raise questions about the feasibility of this process^12,23^. Correct BTB homodimer formation is pivotal to cells, as also highlighted by the dimerization quality control pathway^24–26^ that targets BTB heterodimers and monomers for degradation. BTB domain mutations that impair dimerization are linked to erroneous development and diverse medical conditions^27–29^.

**Figure 1:**
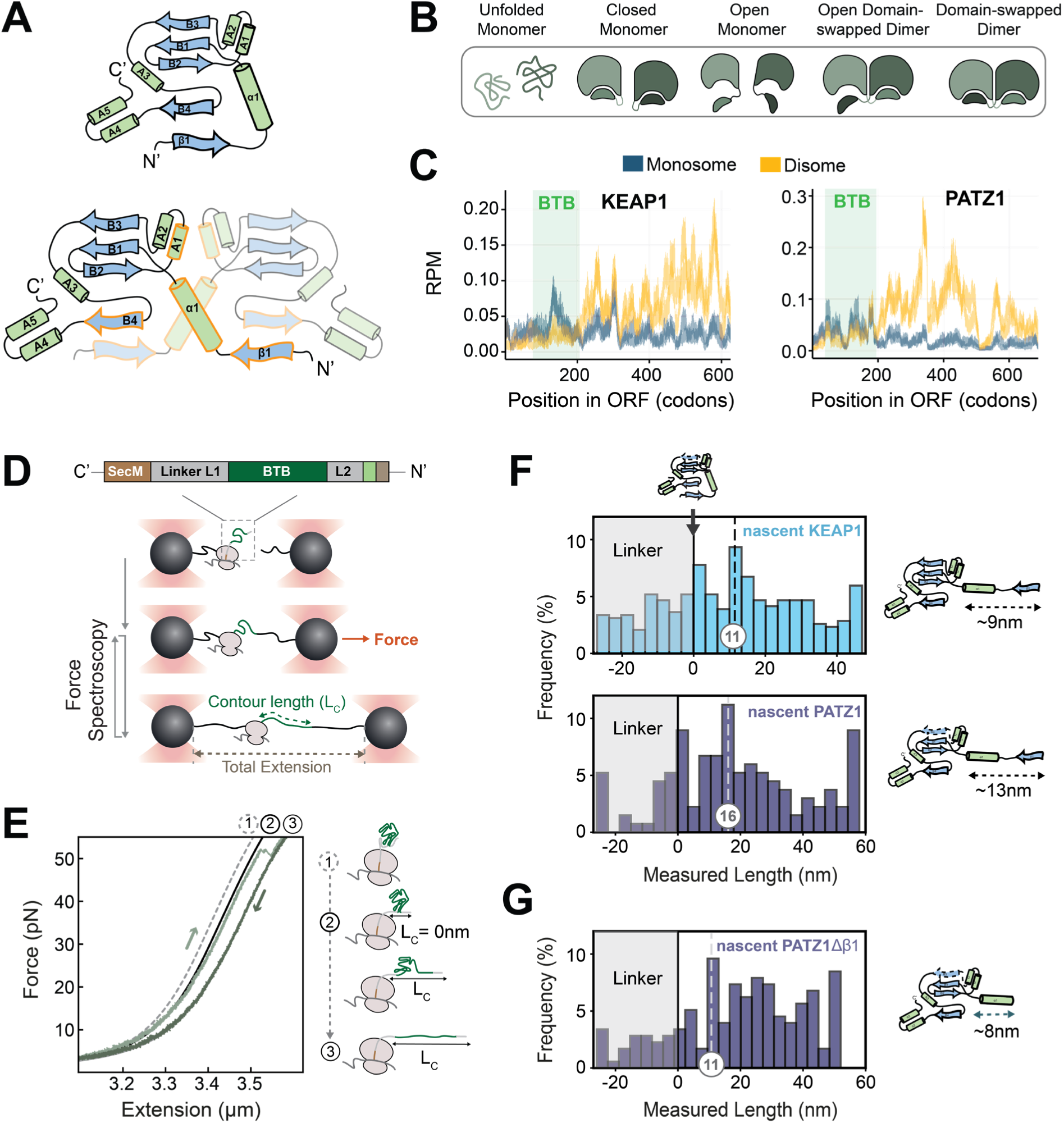
BTB monomers adopt the closed state at the ribosome. A) Topology of closed BTB monomer and BTB dimer. The amino-terminal β-strand β1 folds back onto its own C-terminal β-strand B4 in the closed monomer. In the dimer, β1 folds onto the B4 strand of the other chain. Orange outline: dimerization interface. B) Domain-swapping mechanism cartoon. Closed monomers must open-up to adopt the domain swapped dimer conformation. C) Disome selective profiling (DiSP) for KEAP1 and PATZ1. Monosome (grey) and disome (yellow) footprint density along the position in the ORF (RPM = Reads Per Million). Monosome to disome shift indicated co-co assembly onset. After that disome enrichment levels out and remains high. Data taken for HEK293-T cells^17^. D) Cartoon of optical tweezer approach. SecMstr stalls translation. Flexible Gly/Ser linker L1 pushes BTB domains outside the ribosomal tunnel. Flexible linker L2 allows RNC dimer experiments (see **Fig 3D**, **Fig. S7D**) (see Methods for amino acid sequences). Stalled RNCs are linked to laser-trapped beads by DNA handles (black). Biotinylated nascent chain N-terminus is linked to a DNA handle attached to the second bead by moving the trap and bringing them in closer vicinity. Nascent chains are exposed to repeated stretch-wait-relax cycles by changing the distance between the beads (extension) while measuring the force. E) Example Force-Extension traces for nascent PATZ1 monomer. WLC fit for (1) fully compacted state including linkers, (2) compacted BTB state, (3) fully unfolded state. Deviations from WLC curves indicate changes in unfolded length of the nascent chain. F) Length histogram for nascent KEAP1 (N = 386 states; see methods for further details) and nascent PATZ1 (N = 134 states). Flexible Gly/Ser linkers unfold at low forces in grey area (**Fig. S2B**). Length indicates contour length of the unfolded part of the nascent chain. Fully compacted BTB monomer positions at 0. Dashed line: frequently populated partial fold, consistent with the open monomer state (cartoon right). Cartoons: the N-terminal segment including α1 and β1 is slightly longer for PATZ1 than for KEAP1, estimated as 13 nm and 9 nm, respectively. G) Length histogram for the nascent PATZ1Δβ1 variant (N = 177 states), which consistently shows a lower frequency for the fully compacted closed monomer, and a shift for the open conformation to smaller length.

Here we advance a translation-driven “temporal control” mechanism for the assembly of intertwined protein complexes, and study it at the single-molecule level using BTB dimerization domains as a model system. We hypothesized that translational coupling can offer alternative assembly routes. Specifically, we reasoned that a dimerization domain contains various polypeptide segments that are synthesized in a specific order during translation, and that their ordered emergence could in principle alter the folding-assembly pathway. In our approach, disome selective ribosome profiling (DiSP) is used to detect co-translational assembly of BTB domain proteins *in vivo*, while optical tweezers are employed to study folding and assembly transitions for individual monomers and dimers *in vitro*.

## Results

### *In vivo* dimerization of BTB domain proteins

We assessed BTB dimer formation *in vivo* using Disome Selective Profiling (DiSP) data^17^. In this method, co-co assembly is detected by the conversion of monosomes into disomes during translation, as quantified by deep sequencing of mRNA ribosome footprints, after mRNA digestion and separation of monosome and disomes on a sucrose gradient. We focused on two Kelch proteins: KEAP1, a key sensor of oxidative and electrophilic stress^30–33^ and KLHL12, a substrate-specific adapter of a BCR (BTB-CUL3-RBX1) E3 ubiquitin ligase complex that regulates WNT signaling and ER-Golgi transport^34–36^. We also studied PATZ1, a regulator of embryogenesis, senescence, T-cell development and neurogenesis^37–40^. The DiSP data shows monosome to disome conversion for all these proteins upon translation of their BTB domain (**Fig. 1C, S1A**), thus indicating that their dimerization *in vivo* involves co-co assembly. Structural analysis showed a high degree of structural similarity between the three proteins, each exhibiting a domain-swapped dimer conformation (**Fig. S1B**). PATZ1 contains a 30-amino acid glycine- and alanine-rich central loop between the α-helix A2 and β-strand B3^41^, thus setting it apart from the resolved KEAP1 BTB domain structure^42^ and the AlphaFold 2^43^ predicted structure of the KLHL12 BTB domain.

### BTB monomers adopt a closed state at the ribosome

It is thought that BTB monomers can alternate between a closed state, in which the β-strand β1 is folded back onto the β-strand B4, and an “open” state, in which β1 dissociates from B4 while the rest of the protein remains folded (**Fig. 1B**). The open state is central for BTB dimerization, as it allows β1 to associate with B4 of the dimerization partner. Besides the β1-B4 interaction, the dimerization interface is also composed of the α-helical interactions α1-α1 and A1-A1 (**Fig. 1A**, orange outline). However, native BTB domains are almost exclusively observed as dimers, and attempts to purify BTB monomers have been largely unsuccessful^44–46^, thus leaving the conformation and stability of BTB monomers unclear.

To study monomeric nascent BTB domains we used optical tweezers. We generated stalled ribosome-nascent chain complexes (RNCs) that expose one BTB domain using a modified *in vitro* transcription-translation reaction with biotinylated ribosomes. In a microfluidic chamber, single RNCs were tethered between two laser-trapped polystyrene beads via DNA handles and linked to the ribosome and the N-terminus of the nascent chain (**Fig. 1D**). Nascent chains were probed by repeated stretching and relaxation, and the measured Force-Extension curves were fitted to a worm-like chain (WLC) model (**Fig. 1E**). Sudden or gradual deviations from the WLC model indicated transitions between folded states. Folded states were identified by their contour length, which is the length of the unfolded part of the nascent chain. Consequently, a contour length of 0 nm corresponds to the fully compacted BTB monomer in the closed state (**Fig. 1E**).

We probed KEAP1 and PATZ1 nascent chains during multiple stretch-relax cycles, thus obtaining frequencies of contour lengths indicative of specific folding states (**Fig. 1F**). Both nascent BTB domains showed broad distributions of folding states, with one peak at the fully compacted closed BTB monomer. The entire BTB domain is outside the ribosomal tunnel due to a 42 residue long Glycine-Serine linker at the C-Terminus (**Fig. 1D**). Note that a construct where the BTB domain is pushed outside the tunnel by the natural sequence also showed full compaction (**Fig. S2A**), consistent with the closed monomer state. We observed a second frequently populated contour length that is consistent with unfolding from the closed to the open state. For PATZ1 this folded state had a contour length of 16 nm, slightly longer than the 11 nm observed for the BTB domain of KEAP1 (**Fig. 1F**). Consistently, the N-terminal segment including α1 and β1 is slightly longer for PATZ1 than for KEAP1 (estimated as 13 nm and 9 nm, respectively). To further test whether this folded state is indeed the open conformation, we translated a truncated version of PATZ1 that lacks β1 and hence cannot fold back to form the closed state (**Fig. 1G**). This construct indeed showed reduced frequencies at 0 nm corresponding to the closed state, while the peak for the open state consistently decreased to smaller lengths (from 16 nm to about 11 nm) (**Fig. 1G**). In line with reduced stability for this truncated variant, we further noticed decreased unfolding forces (p = 0.03, **Fig. S2B**) and a wider range of intermediate states (**Fig. 1F-G**). The closed state unfolding force for the full-length constructs was 36 pN and 26 pN for KEAP1 and PATZ1 respectively (**Fig. S2C**). This stability of the closed state, which must open up to dimerize by domain swapping, underscores the challenge of that assembly route.

### BTB closed states resist unfolding

To assess whether ribosome proximity affected BTB domain folding and stability, we performed a set of experiments with full length BTB domains in the absence of ribosomes. We purified KEAP1 and PATZ1 proteins (termed wtKEAP1 and wtPATZ1), as well as the KLHL12 BTB domain (wtKLHL12). To limit measurement noise, short (1.3 kbp) DNA handles were coupled directly to the monomer termini (**Fig. 2A**). We developed a protocol to purify the BTB domains as dimers, attach handles, and subsequently tether BTB domain monomers in between beads (see **Fig. S3**). This method allowed us to obtain stretch-relax cycles on BTB domain monomers (**Fig. 2B**).

**Figure 2:**
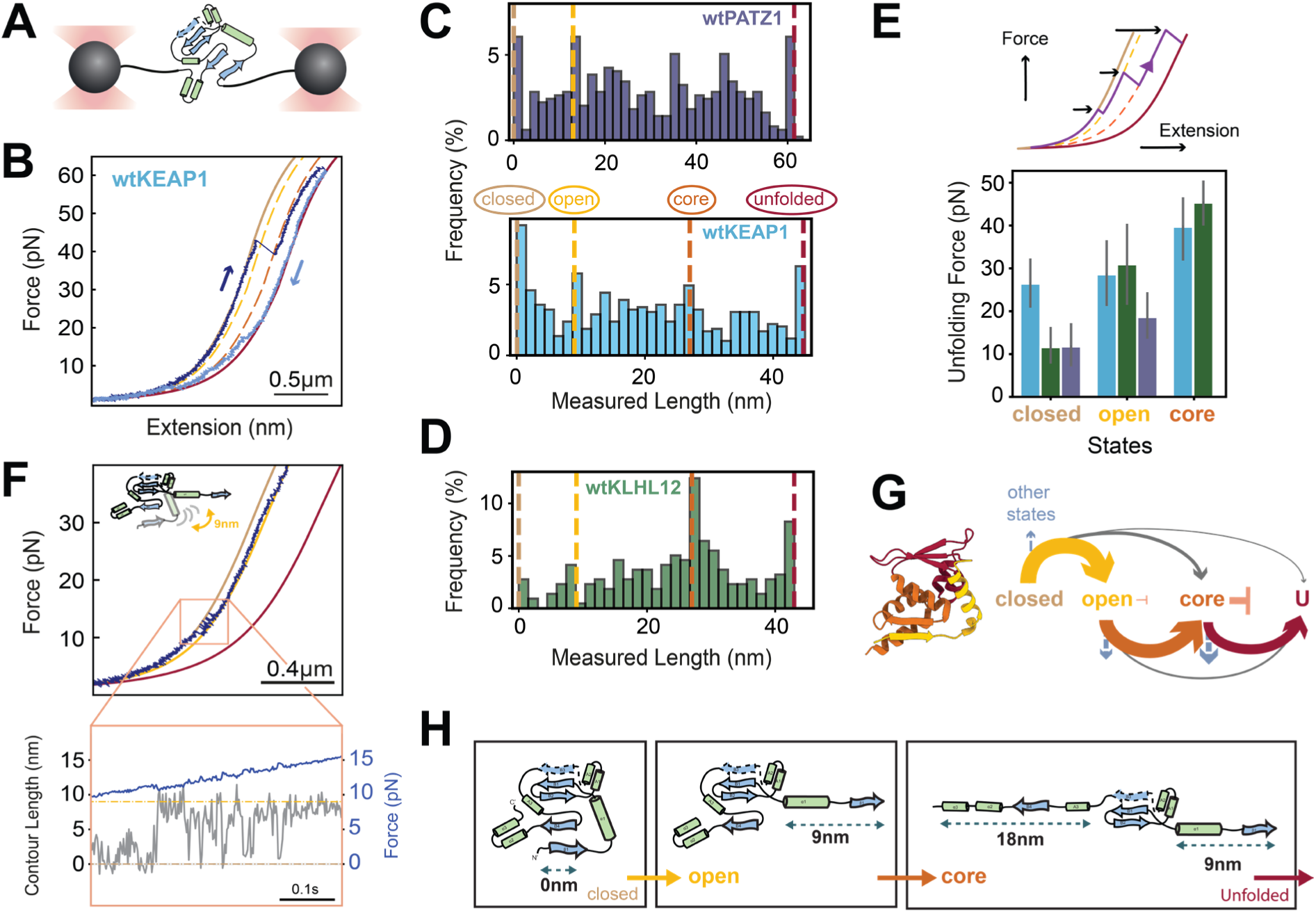
BTB monomers populate a small core fold. A) BTB monomers of KEAP1, PATZ1 and KLHL12 purified and tethered at their C- and N-termini between two optically trapped beads via DNA handles. B) Example force-extension traces for wtKEAP1 monomer, showing unfolding and refolding events indicated by abrupt force changes. C) Contour length histograms showing folded states of wtPATZ1 (N = 495 states; see methods for further details) and wtKEAP1 (N = 589 states). Dotted lines show closed, open, core, and unfolded BTB states. For wtPATZ1 also see **Fig. S5A**. D) Contour length histograms showing folded states of wtKLHL12 (N = 218 states). Dotted lines show closed, open, core, and unfolded BTB states. E) Average unfolding forces of the closed, open and core states, using a length window of 5 nm around the dashed lines. Error bars: 95% confidence interval. Colors: see panels C and D. See methods for N-Values. F) Hopping between “closed” and “open” states (yellow dotted WLC). Zoom: time vs. contour length data showing the hopping behavior, indicating repeated unfolding and refolding between open and closed states. G) Right: Observed transitions between BTB folded states, with thickness indicating frequency (see methods). Left: Structural elements involved in these transitions. H) Unfolding sequence for BTB monomers as derived from the experiments.

Consistent with the nascent BTB domain data (**Fig. 1F**), purified wtKEAP1 and wtPATZ1 populated the fully compacted closed state and the partially folded open state (**Fig. 2C**), with the open state of wtPATZ1 being slightly more extended than that of wtKEAP1. Surprisingly, the unfolding force of the closed state off the ribosome was lower than on the ribosome for wtKEAP1 (26 vs 36 pN p = 5.8·10^−7^), and wtPATZ1 (12 vs 26 pN, p = 1.0·10^−6^) (**Fig. 2E, S2C**). Likewise, the open state also did not show higher unfolding forces for ribosome-released compared to nascent BTB monomers (29 vs 36 pN for wtKEAP1, p = 0.3, and 19 vs 37 pN for wtPATZ1, p = 0.007) (**Fig. 2E**, **S2C**). We observed fast back-and-forth hopping between open and the fully compacted state during stretching (**Fig. 2F**), indicating opening and closing of BTB monomers. Such compaction that can counteract applied forces offers further indication of the stability of the closed state, and the resulting challenges of the domain swapping route.

### BTB monomers populate a small core fold

We wondered if the BTB monomers populate additional folding intermediates, as those could be relevant to monomer-monomer interactions during assembly. Hence, we analyzed intra-chain residue-residue contact maps of the BTB monomers (**Fig. S4**). For KEAP1 and KLHL12, this analysis showed the most contacts in a sub-fold termed the “core” fold, which comprises three β-strands (B1, B2, B3) and two small α-helices (A1 and A2) (**Fig. 2G**, red). Considering that this sub-fold contains roughly 52 residues, it would yield a peak at about 27 nm in the length histograms when it is populated and the rest of the protein is unfolded. Indeed, wtKEAP1 and especially wtKLHL12 showed a pronounced peak at this length within the folded state histogram (**Fig. 2C, D**). Consistently, this core BTB structure also showed the highest unfolding forces, reaching 45.4 pN on average for wtKLHL12 (**Fig. 2E**).

We found that the (un)folding pathways of wtPATZ1 were overall similar to wtKEAP1 and wtKLHL12, but did show minor differences that are consistent with the known structures. Specifically, the residue-residue contact map for PATZ1 showed fewer contacts for this core state, which can be explained by the distinct flexible loop between A2 and B3^41^, and its two rather than three beta-strands (**Fig. S4**). Consistent with these observations, the length histogram showed two instead of one small intermediate (**Fig. 2C, S5A**), which both also showed a lower mean unfolding force (31pN and 33pN) than wtKEAP1and wtKLHL12 (p = 0.01) (**Fig. S5B**).

These data suggest the core and open states as intermediate folded states, that are positioned in between the unfolded state and the closed monomer state, in a manner that is notably similar for all studied proteins. To further test this, we analyzed the transitions between these states (**Fig. 2G**). Consistently, we found that transitions between directly adjacent states along this putative pathway showed the largest frequencies, while transitions that skip one state or go to another state showed lower frequencies (**Fig. 2G, S6A**). Note that the core state, given its significant stability, did not always unfold during stretching to 65pN, which is the maximum because of DNA melting. Refolding was studied by quantifying the state populated after the waiting period at 0 pN to allow for folding without a counteracting force (**Fig. S6B**). This state after folding is most often the fully compacted closed state, while the state before folding is typically the fully unfolded or core state, while the distributions are wide (**Fig. S6B**). Thus, the data indicated the following generic folded states ordered from small to large: core (B1, B2, B3, A1 and A2), open (adding B4, A3, A4, A5), and closed (adding β1 and α1) (**Fig. 2H**).

### Temporal assembly control at the ribosome

Our data suggested the following competing assembly pathways (**Fig. 3A**): After translation of β1, α1 and the BTB core, the BTB cores fold as monomers in both pathways, which split subsequently in two. In the monomer pathway, continued translation exposes B4 that then contacts β1, thus forming the closed monomer that impedes dimerization by domain-swapping (**Fig. 3A**, left). In the dimer pathway, two nascent chains can start interacting after core folding, when helices α1 and A1 of the dimerization interface are exposed but B4 is not (**Fig. 3A**, right). Continued translation exposes the two B4’s that then contact the two β1’s to form the intertwined dimer. This early dimer pathway thus splits off from the monomer pathway before monomer closure is possible, which promotes the direct formation of intertwined dimers (**Fig. 3A**, right).

**Figure 3:**
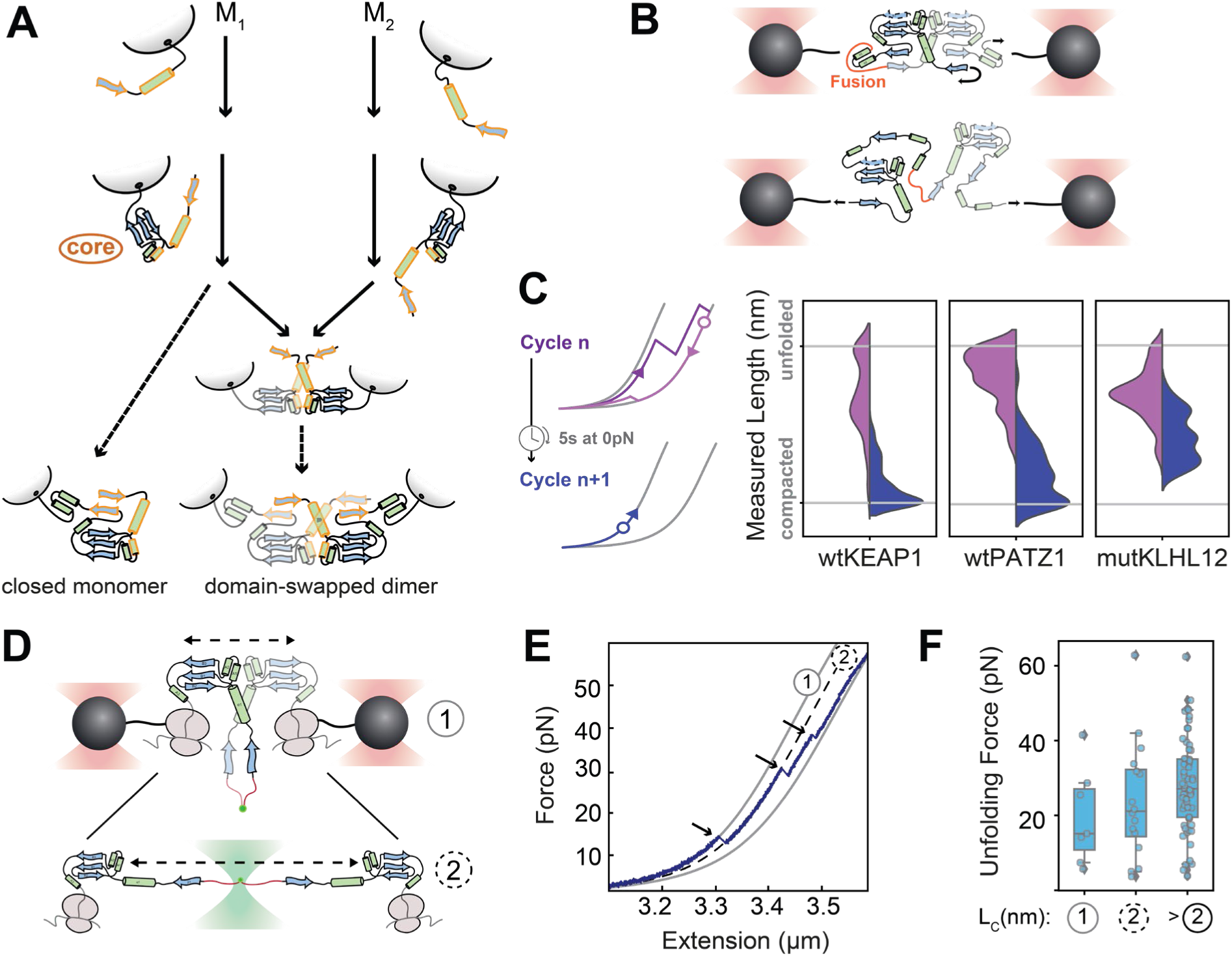
Temporal assembly control at the ribosome. A) Cartoon model for the formation of intertwined BTB dimers using temporal control driven by translation. B) Optical tweezers cartoon for BTB dimer fusion experiments. Fusion^24^ is achieved through a short flexible linker connecting the N-terminus of one subunit chain to the C-terminus of the other subunit chain, which allows for multiple stretch-relax cycles. C) Refolding analysis. Indicated are histograms of the most unfolded state in one stretch-relax cycle (cartoon: purple circle) and the (most compacted) state populated after a waiting period at 0 pN (cartoon: blue circle), for wtPATZ1 (N = 126 events) and wtKEAP1 (N = 234 events), showing a peak at the compacted state indicating dimer formation. In contrast, the dimerization defective mutKLHL12 (N = 136 events) consistently does not show this peak. D) Optical tweezers assay for RNC pair. Two beads are brought together to let nascent chains dimerize. Nascent chains are linked using FlAsH dye^47^, which binds a bipartite tetracysteine motif formed by two N-termini. Dimer interactions between BTB core states (1), which can form state (2) upon dissociation. E) Example force extension trace of an RNC pair (see panel D), showing states sketched in panel D. Dotted lines: matching WLC curves. F) Corresponding unfolding forces around indicated states in panel E within a window of 14 nm around the expected length for (1) and (2). The right-most box shows the forces of unfolding events for unfolded states larger than category (2). N = 7 unfolding events (1), 16 unfolding events (2), 60 unfolding events (3). Compacted states (1) show lower stability against unfolding consistent with the limited dimerization interface that is disrupted upon unfolding or disassembly. Whiskers: 5th and 95th percentiles, box: interquartile range (IQR), center line: median.

To test whether two nearby BTB polypeptide chains interact, and hence form intertwined dimers when allowed to fold, we purified tandem constructs for wtKEAP1 or wtPATZ1 (**Fig. 3B**). In these constructs, two identical BTB monomers were fused head-to-tail, thus exploiting the fact that the N- and C-termini within BTB dimers are close together and hence can be connected by a short linker. The free N- and C-termini of these fused dimers were attached to DNA handles as before. Next, we used optical tweezers to first unfold and then relax them, followed by a 5-10 second time window to refold (**Fig. 3B**). When analyzing the refolds, we observed dominant peaks for the fully compacted state (**Fig. 3C**). These data indicated direct formation of the intertwined dimer state, rather than an initial formation of closed monomers followed by domain swapping. Note that once formed, closed monomers required force to open up (**Fig. 2E, F**), a condition that is not met during the 5-10 s folding time window. To further test dimer formation, we used a dimerization deficient BTB domain of KLHL12 with four mutations in the dimerization interface (L20D/M23D/L26D/L49K)^24^ (**Fig. S1B**). Consistently, here we did not observe the fully compacted state corresponding to the dimer form (**Fig. 3B**). Conversely, a construct based on the wild-type KLHL12 did show the fully compacted dimer state (**Fig. S7A**).

As a final test, we wondered if we could directly observe an early assembly intermediate that pre-empts monomer closure (**Fig. 3A**). This is a challenging aim, as incompletely translated proteins have a strong tendency to aggregate, and hence cannot be purified and interrogated at the single molecule level. In principle, such aggregation can be suppressed in our *in vitro* transcription-translation assay, as the RNCs are attached to beads prior to translation and hence remain spatially separated (**Fig. 1D**). However, these assays probe one RNC while assembly requires two. To address these issues, we developed an assay in which two RNC’s are coupled by linking the N-termini of their nascent chains using a split FlAsH tag^47^ (**Fig. 3D, Fig. S7D**). To focus on the translation timepoint when monomer closure is not yet possible, we arrested translation when B4 (which binds β1) is not yet exposed, while β1, α1 and the BTB core are exposed (**Fig. 3D**). When probing a single RNC as before (**Fig. 1D**), the resulting data were consistent with BTB core refolding (**Fig. S7B, C**). Next, we coupled two RNCs as described above and performed stretch-relax cycles (**Fig. 3D**). We indeed observed full compaction corresponding to an early core-core dimerized state, followed by a length increase consistent with core-core dissociation that produces two tethered monomers in the core state, and finally core unfolding (**Fig. 3E, S7E**). This early core-core dimer state was disrupted at a mean force of 20 pN (**Fig. 3F**). These findings support the suggested model (**Fig. 3A**) in which early assembly intermediates form before monomer closure, which in turn primes the dimer for proper inter-chain β1-B4 docking once B4 is exposed. This assembly pathway uses the order in which nascent chain segments are translated. Specifically, late B4 emergence yields a time window before this B4 emergence in which monomer closure is not yet possible, and the dimerization pathway can be initiated without monomer pathway competition. In this manner, BTB dimers can be formed co-translationally while avoiding the kinetically trapped closed monomer state.

### Assembly control across the BTB proteome

We wondered about the relevance of temporal assembly control for other BTB proteins. Phylogenetic analysis based on BTB domain sequences did not uniformly cluster the entire BTB family, in line with previous work^20^. However, sequences from the BTB-ZF, BTB-BACK-Kelch, and T1 superfamilies did form distinct clusters (**Fig. 4A**, blue, yellow, grey). Other clusters (**Fig. 4A**, black, red, green) were a mix of different superfamilies (e.g. RhoBTB, Skp1, BTB-BACK-Kelch, T1). Consistently, these clusters had broader domain length distributions (**Fig. 4B**). By mapping our DiSP data onto the BTB phylogenetic tree (**Fig. 4A**), we found that the BTB-ZF and BTB-BACK-Kelch superfamilies display a higher frequency of co-co assembly candidates (59% and 70% of the proteins, respectively), compared to the other clusters (**Fig. 4A, C**). Structural analysis using AlphaFold^43^ showed that almost all BTB-ZF and BTB-BACK-Kelch proteins contain the amino-terminal α1, and most also contain β1 (**Fig. S8**). Similarly, almost all co-co assembly candidates contain α1, and most also contain β1 (**Fig. 4D**). These findings agree with our hypothesis that intertwined BTB proteins, which contain α1 and β1, have a higher need for the temporal control that co-co assembly enables. Consistently, BTB domain proteins from the T1 superfamily proteins all lack α1 and β1 (**Fig. S6**) – and none of them is a high-confidence co-co candidate (**Fig. 4A, C, D**).

**Figure 4:**
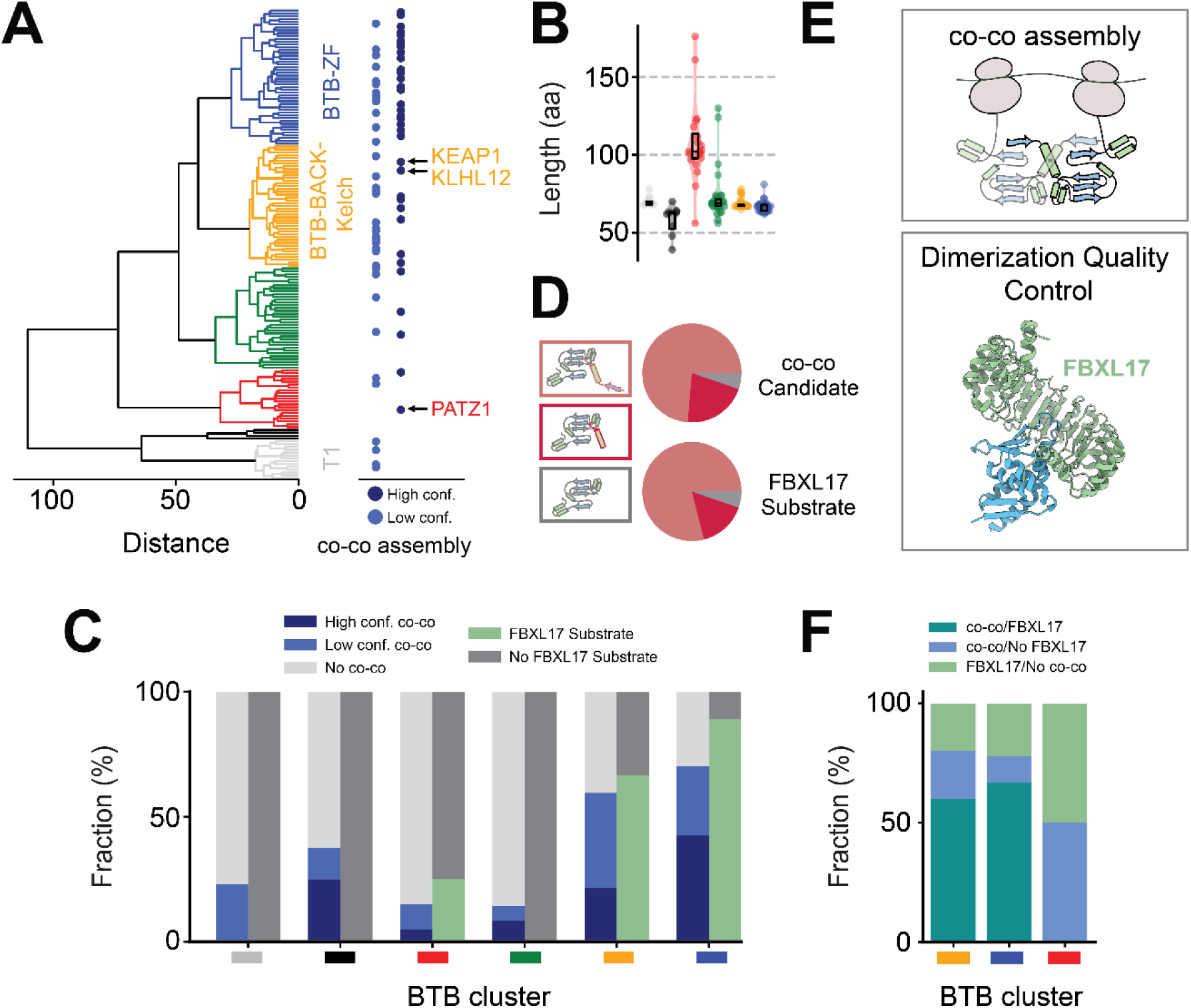
Assembly control across the BTB proteome. A) Phylogeny based on 166 BTB domain sequences (Uniprot^64^). Colors indicate different BTB sequence clusters. High (N = 36) and low (N = 37) confidence coco-assembly proteins^17^ are highlighted by light and dark blue dots respectively. B) Length distribution of for BTB domains as annotated in Uniprot for the different clusters. Clusters which could not be assigned to one BTB superfamily had a larger spread of lengths (box: standard deviation) compared to the superfamily clusters of the BTB-ZF, BTB-BACK-Kelch and T1 superfamilies (box: standard deviation). C) Fraction of co-co assembly candidates^17^ and FBXL17 substrates^25^ for the different sequence clusters. As FBXL17 substrates were not tested genome-wide the fractions were normalized to the total amount of tested substrates per cluster. D) Structural analysis of the BTB amino-terminal extension. AlphaFold^43^ structure predictions show that BTB co-co assembly candidates^17^ and FBXL17 BTB substrates^25^ predominantly either contain the full amino-terminal extension or only α1. E) Two assembly control mechanisms. Top: Translational coupling enables the intertwining of polypeptides for BTB dimer formation. Bottom: Aberrant BTB dimers or monomers (shown here) are recognized by SCF^FBXL17^ (pdb: 6w66), a dimerization-quality-control E3 ligase, which ubiquitylates its substrates for degradation. F) Comparison between FBXL17 substrates and co-co assembly candidates shows that most FBXL17 substrates are also co-co assembly candidates.

The importance of BTB assembly control mechanisms is highlighted by the recently discovered dimerization quality control pathway, in which SCF^FBXL17^ ubiquitinates and helps to degrade aberrant BTB heterodimers and monomers during and after translation (**Fig. 4E**). FBXL17 is thought to detect aberrant dimers by differences in the stability compared to native dimers^24–26^. The BTB-ZF and BTB-BACK-Kelch BTB superfamilies utilize co-co assembly most extensively (**Fig. 4C-D**), which could indicate they are the most challenging to assemble and hence are more frequently FBXL17 substrates. A similar mapping showed that this is indeed the case (**Fig. 4C**). In addition, almost all FBXL17 substrates^25^ contain the α1 and β1 amino-terminal extension (**Fig. 4D**). Consistently, within the BTB-ZF and BTB-BACK-Kelch superfamilies, the majority of FBXL17 substrates have also been detected as co-co assembling candidates (**Fig. 4F**). Note that the FBXL17 substrate test did not cover the complete genome, hence for many co-co assembly candidates it is unknown if they are also FBXL17 substrates. Thus, the correlation between co-co assembly and FBXL17 interaction may be even stronger. Overall, these analyses are consistent with the idea that intertwined BTB dimers benefit from assembly control mechanisms, as offered by translation-driven temporal assembly control and FBXL dimerization quality control^25^.

## Discussion

While many protein complexes show intertwined subunits, their formation remains poorly understood. The conformational monomer remodeling required for their assembly suggests major activation energies and assembly delays^23,48–51^, which in turn poses aggregation risks^52^. Here, we advanced a temporal control mechanism driven by translation that circumvents this issue (**Fig. 3A**). It avoids remodeling requirements by preventing unproductive conformations in the first place. Interactions between nascent chains and their progressive emergence during translation are central. Specifically, the mechanism exploits the fact that newly emerged chain segments are transiently safeguarded against unproductive contacts with soon-to-emerge segments encoded upstream, offering an opportunity to form alternative productive contacts with another nascent chain – even if the latter have low probability to form normally, during later phases or after translation. Assembly-folding processes hence can follow pathways that are poorly accessible without temporal control. Hence, by using the energy-driven nature of translation, these assembly-folding pathways break the constraints of equilibrium assembly thermodynamics, to avoid unproductive intermediates directed towards native dimeric states.

As shown for KEAP1, KLHL12 and PATZ1, BTB domain monomers risk forming unproductive stable closed states, in which one β-strand (β1) folds back on another (B4), at the ribosome and away from it (**Fig. 3A**, left). During translation however, these intra-chain contacts are excluded until B4 emerges, while early inter-chain contacts can already form (**Fig. 3A**, middle). The β1 segments are then well positioned to fold onto the subsequently emerging B4 of the partner subunit, completing the intertwined dimer structure. The mechanism makes use of the sequential emergence of nascent chain segments to promote certain interactions over others, thus steering folding-assembly pathways away from closed BTB monomers.

Temporal assembly control offers an alternative to domain swapping, which has been studied extensively for several proteins including BTB domains^20^. Domain swapping is non-trivial to study, given that domain-swapped dimers interconvert slowly^48,49^, and closed BTB monomers have been difficult to detect^44–46^, possibly due to hinge loop conformations and other stabilizing interactions between subunits^45,48^. Our approach enabled BTB monomer studies, and revealed a high stability of closed KEAP1, KLHL12 and PATZ1 monomers, as evidenced by substantial unfolding forces (10 – 35 pN) (**Fig. 2E, S5B**). Stable closed monomers were also readily formed co-translationally in absence of dimerization partners (**Fig. 1F, Fig. S2A,C**), despite reported destabilizing influence of ribosomes on nascent folds^53^. These stable closed BTB monomers are a barrier for efficient dimerization, and hence underscores the importance of alternative assembly pathways as advanced here.

To explore *in vivo* and cross-proteome relevance we analyzed BTB phylogeny and structural features, and correlated these with our single-molecule and disome selective ribosome profiling data. While the PATZ1 BTB domain we studied at the molecular level has significant phylogenetic distance to KEAP1 and KLHL12, we found that their folding intermediates and assembly mechanisms were notably similar (**Fig. 1-3, Fig. S5**), with small differences in unfolding step sizes consistent with the longer N-terminal extension and the additional flexible loop within the core fold of PATZ1 (**Fig. 1F, 2C, S5**). The analysis showed co-co assembly and FBXL17 interactions are most prevalent in the BTB-BACK-Kelch and BTB-ZF superfamilies (**Fig. 4A, E**), with β1 and α1 in the amino-terminal extension being the main structural determinants (**Fig. 4D**), which is consistent as they contribute to dimerization. Interestingly however, a certain fraction of BTB domains that lack β1 within the Alpha-Fold predictions do display co-co assembly and FBXL17 interactions (**Fig. 4D**). This observation is in line with the notion that dimerization can commence without β1-B4 interactions, before B4 emerges (**Fig. 3A**). An example is MIZ1, which lacks β1 but does form a homodimer ^45^.

While the DiSP method could show prevalence^17^, it could not address the functional relevance of co-co assembly. Here we show that co-co assembly opens up alternative folding-assembly pathways that avoid kinetically trapped states and slow domain-swapping^48,49^. The involved temporal control provides a mechanism to assemble intertwined dimers that are challenging to form otherwise, and hence expands the spectrum of protein structures that can be faithfully synthesized, as well as limiting unwanted complexes such as BTB heterodimers^54^. The emergence of polypeptide segments in a specific temporal order is general for all proteins.

Temporal assembly control may thus be exploited more generally to enable complex protein structures beyond BTB domains. Our results indicate that assembly-folding pathways can be subjected to regulatory control, which can involve evolutionary tuning of various functions. For instance, assembly pathways may be shaped by processes that affect ribosome-ribosome proximity^55,56^, including translation speed and pausing^57–59^, direct ribosome-ribosome interactions, as well as by protein folding topologies^60–63^. Our findings further suggest that temporal assembly control works in conjunction with dimerization quality control systems such as FBXL17, which act both co- and post-translationally. This interplay may be reciprocal, with its quality control factors regulating the assembly control process or rescuing unsuccessful dimerization.

## Acknowledgements

Work in the group of S.J.T. is supported by the Netherlands Organization for Scientific Research (NWO). M.B. and K.F. were supported by a HBIGS PhD fellowship. M.B. was additionally supported by a Boehringer Ingelheim Fonds (BIF) PhD fellowship. B.B. and S.J.T. acknowledge a research grant of the European Union (ERC - SyG - 101072047 - CoTransComplex). Views and opinions expressed are however those of the authors only and do not necessarily reflect those of the European Union or the European Research Council. Neither the European Union nor the granting authority can be held responsible for them.

## Data availability

Sequencing data reported in this study is available at GEO under accession number GSE151959.

## Software availability

Data analysis of ribosome profiling datasets were performed with RiboSeqTools^17^ (available at https://github.com/ilia-kats/RiboSeqTools). Other Python code and data is available at Zenodo/Github (https://doi.org/10.5281/zenodo.15396113).

## Author Contributions

Conceptualization: K.T., V.S., M.B., P.J., F.T., B.B., G.K., M.R and S.J.T. Methodology: K.T., V.S., P.J., M.B., F.T., J.S., M.R., B.B., G.K., S.J.T. Biotinylated ribosomes: A.K. Disome Selective Profiling Experiments: M.B. and K.F. Single-molecule experiments: K.T. Single molecule data analysis and visualization: K.T., S.J.T. Writing: K.T. and S.J.T. Supervision: S.J.T., M.R., B.B. and G.K.

## Methods

### Disome Selective Profiling (DiSP)

We grew U2OS (ATCC Cat# HTB-96, RRID: CVCL_0042) and HEK293-T cells (DSMZ Cat# ACC 635) in high glucose DMEM media containing GlutaMAX and pyruvate (Gibco) with 10% heat-inactivated FCS (Gibco), 100 units/mL penicillin and 100 μg/mL streptomycin (Gibco) and in a humidified incubator with 5% CO2 at 37°C (HERAcell 150i). For DiSP of HEK293-T and U2OS cells, lysis buffer contained a physiological salt concentration (50 mM HEPES pH 7.0, 10 mM MgCl_2_, 150 mM KCl, 1% NP40, 10 mM DTT, 100 μg/ml CHX, 25 U/ml recombinant Dnase1 (Roche) and protease inhibitor (complete EDTA free, Roche). For DiSP of HEK293-T cells, we used a high-salt lysis buffer containing 500 mM KCl for possible effects on the onset of disome formation. For DiSP of U2OS cells, lysis was performed with chemical crosslinkers, using 2.5 mM BS3 and 20 mM EDC. For crosslinking to occur with cell lysis, cells were scraped in crosslinker-containing lysis buffer on ice. After lysis, DiSP samples were processed^1^. Briefly, we clarified cell lysates, loaded them on sucrose gradients (5% - 45%), centrifuged them for 3.5 hours at 35,000 rpm, 4°C (SW40-rotor, Sorvall Discovery 100SE Ultracentrifuge) and collected fractions corresponding to monosomes and disomes. We extracted 30 nt long footprints from both fractions and deep sequenced them. In the DiSP gene density profiles, we show the position-wise 95% Poisson confidence interval corrected for library size, smoothing the read counts with a 15-codon wide sliding window^1^.

### Cloning

For in vitro transcription, the SecMsrt site was added on C-Terminus of pRSET vector (Thermo Fisher). PATZ1 and KEAP1 were cloned using NdeI and SpeI (New England BioLabs) restriction enzymes. The Plasmid DNA of PATZ1 was kindly obtained from the Bukau lab at the Center for Molecular Biology of Heidelberg University (ZMBH) and the DNA plasmids of KEAP1 and KLHL12 from Michael Rapé Lab at the UC Berkley and the Howard Hughes Medical Institute (HHMI). DNA of the gene of interest was amplified by PCR using a gene specific primers and Phire Green Master mix (Thermo Fisher). The fragments were purified from the gel with QIAquick gel purification kit (Qiagen) and digested with restriction enzymes (New England BioLabs), followed by purification with QIAquick PCR purification Kit (Qiagen). 30ng of digested vector and appropriate amount of insert (1:5 ratio) were ligated using 400U of Hi-T4 ligase (#M2622S, New England BioLabs) in 20 µl set up at room temperature for 1 hour. Ligation mix was used for a heat shock transformation in Dh5-alpha competent cells (New England BioLabs). The cells were allowed to recover at 37°C for up to 1 hour, followed by plating on the LB plate supplemented with 100 µg/ml ampicillin and grown overnight at 37°C. Subsequently, positive colonies were grown again in LB medium supplemented with 100 µg/ml ampicillin and used for a plasmid isolation (QIAprep Spin Miniprep Kit, Qiagen). The sequences were verified by sequencing (Eurofins, Germany).

#### Nascent Chain Constructs: Amino Acid Sequences

<u>1351: pRSET-amber-FlAsH-14aaLinkerL2-**Keap1(50-179)-**42aaLinkerL1- secMsrt</u>

M*GGCCEGKSSGSGSESKSTSMGS**MRTFSYTLEDHTKQAFGIMNELRLSQQLCDVTLQVKYQDAPAAQFMAHKVV LASSSPVFKAMFTNGLREQGMEVVSIEGIHPKVMERLIEFAYTASISMGEKCVLHVMNGAVMYQIDSVVRACSDFLV QQ**GTEGKSSGSGSESKSTEGKSSGSGSESKSTEGKSSGSGSESKSTTSFSTPVWIWWWPRIRGPP

<u>1340: pRSET-amber-FlAsH-14aaLinkerL2-**Keap1(50-158)-**secMsrt</u>

M*GGCCEGKSSGSGSESKSTGST**MRTFSYTLEDHTKQAFGIMNELRLSQQLCDVTLQVKYQDAPAAQFMAHKVVL ASSSPVFKAMFTNGLREQGMEVVSIEGIHPKVMERLIEFAYTASISMGEKCVLHVMN**GTSFSTPVWIWWWPRIRGPP

<u>1370: pRSET-amber -**Keap1(50-197)-**secMsrt</u>

M*T**MRTFSYTLEDHTKQAFGIMNELRLSQQLCDVTLQVKYQDAPAAQFMAHKVVLASSSPVFKAMFTNGLREQGM EVVSIEGIHPKVMERLIEFAYTASISMGEKCVLHVMNGAVMYQIDSVVRACSDFLVQQLDPSNAIGIANFAEQIGCV**TS FSTPVWIWWWPRIRGPP

<u>1210: pRSET-amber-FlAsH-14aaLinkerL2-**Patz1(1-166)-**42aaLinkerL1-secMsrt</u>

M*GGCCEGKSSGSGSESKSTSMG**ERVNDASCGPSGCYTYQVSRHSTEMLHNLNQQRKNGGRFCDVLLRVGDESF PAHRAVLAACSEYFESVFSAQLGDGGAADGGPADVGGATAAPGGGAGGSRELEMHTISSKVFGDILDFAYTSRIVV RLESFPELMTAAKFLLMRSVIEICQEVIKQSNVQILVP**GTEGKSSGSGSESKSTEGKSSGSGSESKSTEGKSSGSGSE SKSTTSFSTPVWIWWWPRIRGPP

<u>1211: pRSET-amber-FlAsH-14aaLinkerL2-**Patz1(18-166)-**42aaLinkerL1-secMsrt</u>

M*GGCCEGKSSGSGSESKSTSM**VSRHSTEMLHNLNQQRKNGGRFCDVLLRVGDESFPAHRAVLAACSEYFESVFS AQLGDGGAADGGPADVGGATAAPGGGAGGSRELEMHTISSKVFGDILDFAYTSRIVVRLESFPELMTAAKFLLMRS VIEICQEVIKQSNVQILVP**GTEGKSSGSGSESKSTEGKSSGSGSESKSTEGKSSGSGSESKSTTSFSTPVWIWWWPRI RGPP

Star (*) denotes the UAG (amber) codon that allows introduction of a biotin moiety with a triple aminophenyl spacer.

### Coupling of ribosomes to beads with DNA handles

5kbp long double-stranded DNA (dsDNA) molecules were prepared by PCR amplification using digoxigenin (DIG) and biotin 5’-end-modified primers. Neutravidin (NTV) (ThermoFisher, 31000) was added in a 200 times excess ratio to the PCR fragments (Bio-DNA-DIG) and incubated overnight at 4°C in a rotary mixer.

Two batches with 0.14mg/ml anti-digoxigenin coated polystyrene beads (Speherotech, DIGP-20-2) with a diameter of 2.1 μm were incubated in 12 μl TICO buffer (20 mM HEPES-KOH pH 7.6, 10 mM (Ac)_2_Mg, 30 mM AcNH_4_, 4 mM β-mercaptoethanol as an additional oxygen scavenger) with 1.4 nM NTV-Bio-DNA-DIG handles at 4°C in a rotary mixer for around 30 minutes. Afterwards unbound DNA was removed by pelleting and washing NTV-Bio-DNA-DIG coupled beads twice by centrifugation at 4°C at 3000 rpm for 5 minutes and resuspending in fresh TICO buffer each time. After the last washing step one bead batch was resuspended in 20 μl of TICO buffer and the other in 250 μl. Within the 20 μl, 1.2 U/μl RNase Inhibitor, Murine (New England BioLabs, M0314S) is added together with 390 nM ribosomes that were biotinylated in vivo at the uL4 ribosomal protein on the large subunit and subsequently isolated as previously described^2^ from Can20/12E^3^. The mixture was incubated at 4°C in a rotary mixer for around 45 minutes. Excess unbound ribosomes were removed by two consecutive washing steps with TICO buffer. After the last washing step the mixture was directly resuspended in a modified *in vitro* transcription-translation reaction to generate stalled ribosome-nascent chain complexes (RNCs) as described below.

### Cell-free protein synthesis

RNCs of PATZ1 and KEAP1 BTB domains were generated through an *in vitro* transcription-translation reaction using a modified version of the PURE system^4^ lacking ribosomes (New England BioLabs, E3313S). The system was supplemented with 10 μM of modified tRNA pre-charged with biotinylated lysine (Bio-Connect, PRX-CLD04) to co-translationally incorporate a biotin-tag at the N-terminus of the nascent chain with an amber stop codon and 0.83U/μl RNase Inhibitor, Murine (New England BioLabs, M0314S). 5.5 nM linearized plasmid was added to the reaction mixture after mixing it with the biotinylated ribosome bound beads. Synthesis was carried out at 37°C for 20 min. The bead-tethered RNCs were resuspended in 300 μl TICO buffer and injected in the microfluidic chamber.

### Preparation of the FlAsH Dye

FlAsH-EDT₂ (Carbosynth) stock solution was prepared at a concentration of 5 mM by dissolving in DMSO (Thermo Scientific) and storing it at −20 °C under an inert atmosphere^5^. For experiments, the stock solutions were diluted in TICO buffer to a working concentration of 500 nM.

### Protein expression and purification

The ybbr tag (DSLEFIASKLA) was added on both termini of KEAP1, KLHL12 and PATZ1 proteins.^6^ The proteins were fused to MBP (maltose binding protein). The fused proteins were cloned into pet28(a) plasmid (Novagen) and transformed into Rosetta strain (gift from Matthias Mayer, Germany). The cells were grown in LB medium supplemented with 50 µg/ml kanamycin until an OD of ∼0.6 was reached. Expression was induced by adding 0.2 µM IPTG (Sigma) at 15°C overnight with slow agitation. Cells were harvested by centrifugation at 5000rpm (Beckman) for 20 min. The pellet was resuspended in a pre-chilled lysis buffer (50 mM phosphate buffer, pH 7.5, 200 mM NaCl, 10 mM EDTA, 50 mM Glutamic Acid–Arginine, 1 mM DTT (Sigma) and protease inhibitor (cOmplete^™^ Mini, EDTA-free (Roche) and lysed using an Emulsiflex homogenizer. To remove insoluble material, centrifugation at 50,000 g for 1 hour at 4°C was performed. The clarified lysate was incubated with Amylose resin (NEB) for 1 hour at 4°C. After washing the resin extensively with buffer A (50 mM Tris-HCl, pH 7.5, 200 mM NaCl, 1 mM DTT), the bound proteins were eluted with buffer A containing 20 mM maltose. Lastly, maltose was removed in a PD10 desalting column (GE Healthcare) and eluted in SFP coupling buffer (50mM Hepes, 10 mM MgCl₂, pH 7.4).

### Attachment of DNA handles to purified BTB monomers and dimers and to beads

The purified BTB proteins were conjugated to a 20 nt oligo modified with coenzyme A (Biomers GmbH) in the presence of 50 mM HEPES, 10 mM MgCl₂ (both from Merck), and 1 µM SFP synthase (Addgene) at 4°C overnight. Any remaining oligos were removed through Ni-NTA purification (Protino, MACHEREY-NAGEL).^6^

To create the 1.3 kb DNA handles, a PCR amplification was performed using primers with phosphate at one end and either biotin or digoxigenin at the other end (Eurofins, Germany), starting from the pUC19 plasmid (New England Biolabs). The amplified DNA was purified using the Qiagen PCR Purification Kit (Qiagen). The DNA was then digested with Lambda exonuclease (New England Biolabs) at 37°C for 2 hours, followed by 10 minutes of inactivation at 75°C.

An equal amount of both ssDNAs (biotin and digoxigenin) were mixed and purified together using an Amicon 30 kDa column (Merck, Darmstadt, Germany). The second strand was completed using Deep Vent (exo-) DNA polymerase (New England Biolabs), dNTP mix (ThermoFisher), and a phosphorylated primer (Eurofins, Germany), starting 20 nt downstream to generate the overhang. The final product was purified using the Monarch PCR Purification Kit.

The 1.3 kb DNA handles with a 20 nt overhang complementary to the oligo attached to the protein, were ligated to the protein-oligo complex using T4 ligase (New England Biolabs) at 16°C for 4 hours, followed by an overnight incubation on ice. The resulting DNA-BTB-DNA complex were either stored on ice for a few days, if directly measured, or flash-frozen and stored at −80°C for future analysis.

Before measurement the DNA-BTB-DNA construct was mixed with anti-digoxigenin coated polystyrene beads (Spherotech, DIGP-20-2) and incubated together at 4°C in a rotary mixer for around 30 minutes.

### Optical tweezers assay and single-molecule data analysis

Data was recorded using a C-Trap instrument (Lumicks, Amsterdam) equipped with a single high-intensity, polarization-stable 1064 nm laser. The laser is split into two orthogonally polarized beams, one of which can be steered using a piezo mirror relative to the other, enabling dual optical trapping. Additionally, the system includes two fluorescence excitation lasers (532 nm and 638 nm) for multi-color confocal fluorescence detection. This allows to carry out correlated single-molecule force spectroscopy and multi-color confocal laser scanning spectroscopy. Single-photon sensitivity is achieved through the use of photodiodes (APDs). Calibration of the optical traps was performed using the power spectrum method^7^, where the power spectra of trapped beads undergoing Brownian motion were fitted with a Lorentzian function, obtaining average stiffness values of k = 0.35 ±0 .045 pN nm^-1^.

Measurements were conducted in a monolithic laminar flow cell equipped with a pressure-driven microfluidic system featuring five separate flow channels. This design ensures the separation of beads carrying ribosome-nascent-chain complexes (RNCs) or purified protein from bead-tethered NTV-Bio-DNA-DIG constructs or NTV coated beads (Spherotech, NVP-20-5), respectively.

During experiments, individual molecules were tethered by trapping a bead from each flow channel and transferring them into a separate measurement side channel. This channel was supplemented with a P2O oxygen scavenging system containing 3 U/ml pyranose oxidase, 90 U/ml catalase, and 50mM glucose (Sigma) to reduce damage by reactive oxygen species induced by the trapping laser and to keep the pH stable. To form a tether of individual molecules, optically trapped beads were brought within close proximity, such that either the biotin tag at the end of the RNC constructs could link up to the NTV-Bio-DNA-DIG constructs attached to the other bead or in the case of the purified BTB proteins bound to the anti-DIG coated beads, the free biotin-DNA handle could bind to a NTV coated bead (Fig. S3). Tethers other than from individual molecules may form between the optically trapped beads, which are not aligned in parallel with the bead-bead axis resulting in a shorter bead-bead distance, do not follow the WLC model, do not show proper unfolding, do not yield a single step break when one tether breaks, and can hence be identified. Measurements were conducted using a cycling force spectroscopy mode, in which the steerable optical trap was moved at a constant velocity of 0.1 μm/s and a maximum force of up to 65 pN. The resulting force-extension curves for individual tethers were analyzed with a custom written python script by fitting two worm-like chain (WLC) models in series: a twistable worm-like chain (tWLC) model to account for the DNA handle contribution^8^ and an inextensible WLC model for the protein contribution^9^, yielding the following relation between extension (x) and force (F) is given:

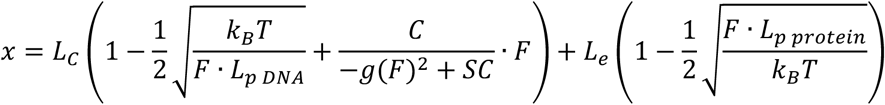

The first term parameterizes the DNA by: the contour length *L*_c_ (884nm for 1.3kb and 3400nm for 5kb DNA handles), the twist rigidity *C* (literature^10^: 440±40pN nm^2^), the stretching modulus *S* (863pN/nm (SD 183pN/nm)), the persistence length of the DNA *L*_*p* *DNA*_ (average values: 37nm (SD of 10 nm) and the twist–stretch coupling *g*(*F*). The second term is the inextensible WLC model for protein contribution, where *L*_*p protein*_ (0.75nm) and *L*_e_ are the persistence and extended length of the protein, respectively. N-values Fig. 1F: KEAP1 (N = 7 molecules, 53 cycles), PATZ1 (N=8 molecules, 34 cycles). N-values Fig. 1G: PATZ1 Δβ1 variant (N = 7 molecules, 25 cycles). N-values Fig. 2C: wtPATZ1 (N = 7 molecules, 72 cycles), wtKEAP1 (N = 10 molecules, 123 cycles). N-values Fig. 2D: wtKLHL12 (N = 5 molecules, 36 cycles). N-Values Fig. 2E for bars from left to right, respectively: 53, 7, 23, 37, 15, 39, 26, and 32 unfolding events. N-values Fig. 3C: wtPATZ1 (N=5 molecules, 68 cycles), wtKEAP1 (N=9 molecules, 127 cycles) and mutKLHL12 (N = 5 molecules, 73 cycles). N-values Fig. 3F: 7 (states 1), 16 (states 2), 60 (states 3) unfolding events. N-values Fig. S2B for bars from left to right, respectively: 86, 18, 22, 129, 57, 77 unfolding events. N-values Fig. S2C for bars from left to right, respectively: 22, 8, 27, 11 unfolding events. N-values Fig. S5A: wtPATZ1 (N = 7 molecules, 72 cycles). N-Values Fig. S5B for bars from left to right, respectively: 53, 7, 23, 37, 15, 39, 26, 32, 20, and 27 unfolding events. N-Values Fig. S5C: number of cycles initially in closed state: 22, in open state: 38, in core2 state: 28, in core3 state: 41. N-values Fig. S6A: number of cycles initially in closed state: 47 (wtKeap1), 8 (wtKLHL12), in open state: 41 (wtKeap1), 14 (wtKLHL12), and in core state: 48 (wtKeap1), 27 (wtKLHL12). N-values Fig. S7A: wtKLHL12 (N = 2 molecules, 28 cycles). N-values Fig. S7C: KEAP1 (50-159aa) (N=15 molecules, 38 cycles). N-values Fig. S7E: KEAP1 (N = 13 molecules, 31 cycles). The Mann-Whitney U test was used for statistical testing to obtain the p-value.

### Calculation of transition probabilities

A molecule was classified as being in the closed, open, core, or unfolded state if its proposed length fell within a ±3 nm range around the length of the respective state. Based on this classification, the cycles of individual molecules, which transitioned to a state, with a larger length during stretching, or any other state of different length, or remained in the same state without length changes, were counted. Transitions may occur via an intermediate state (e.g., from open to an intermediate state between open and core before reaching core). Transition frequencies were calculated by dividing the number of cycles with a specific transition by the total number of cycles, which included one of the defined states marked as a starting point for each specific transition (see **Fig. S6A**).

### Hierarchical clustering of BTB domain sequences

BTB domain positions and sequence were obtained from Uniprot using only reviewed SwissProt entries. BTB domains were labeled with high or low confidence assembly if at least 30 residues of the BTB domain were emerged from the ribosomal exit tunnel at the time of coco-onset as determined in Bertolini et. al.^1^ Table S1. Pair-wise sequence alignment scores of all BTB domains were obtained using the biopython pairwise2.align.globaldx module with blosum62 alignment matrix. Finally, scores were hierarchically clustered using the scipy linkage module and the ward method.

## Supplementary Figures

**Supplementary Figure 1.**
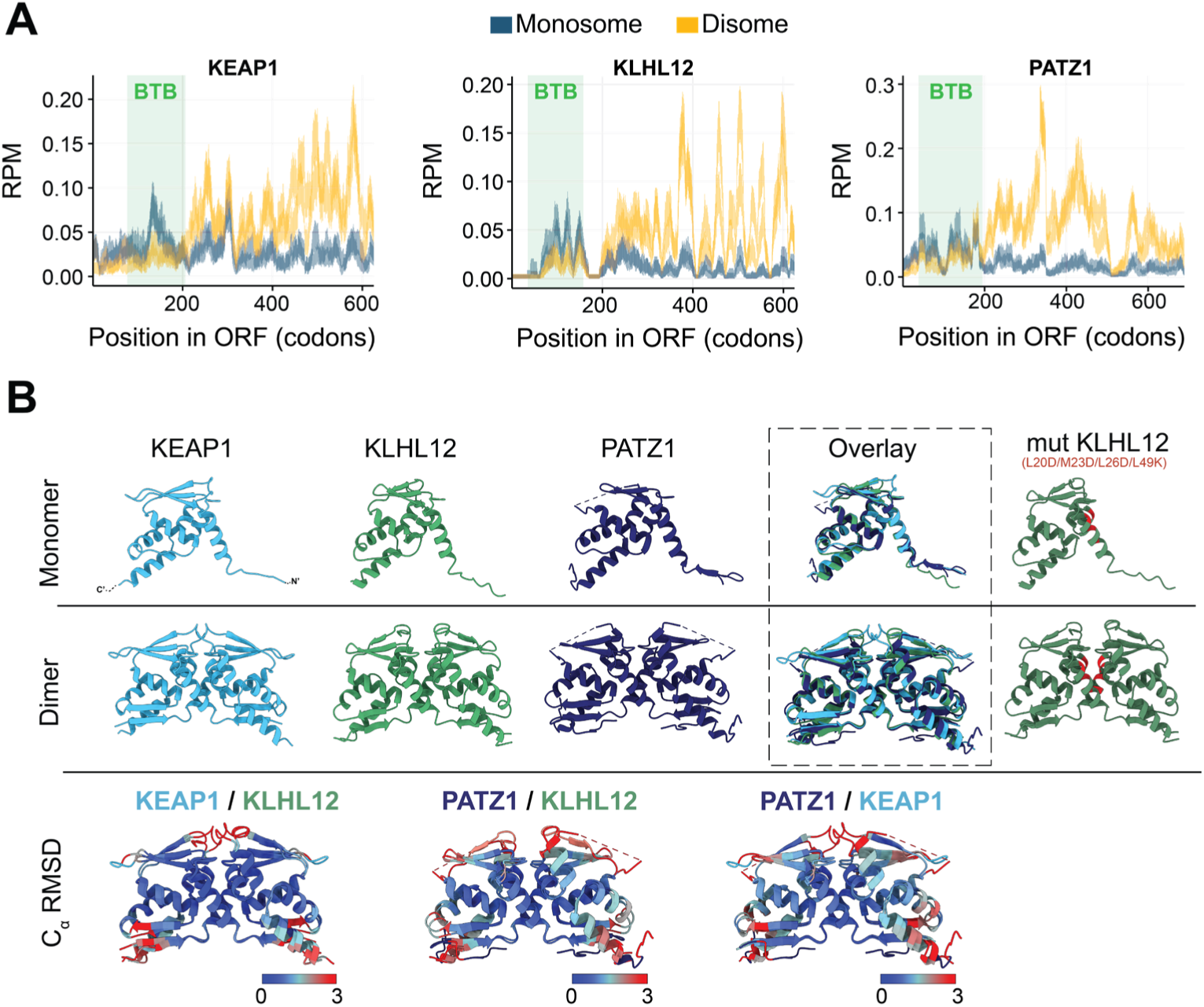
A) Monosome (grey) and disome (yellow) footprint density along the position in the ORF (RPM = Reads Per Million) of KEAP1, KLHL12 and PATZ1. Green bars indicate the position of the BTB domain. A monosome to disome shift of translating ribosomes, the onset of co-co assembly, is observed after the exposure of around 200 amino acids outside of the ribosomal tunnel, shortly after the BTB domain. After that disome enrichment levels out and remains high. DiSP data is from HEK293-T cells^1^. B) Structures of KEAP1 (pdb: 7EXI), KLHL12 (AlphaFold^2^ prediction: AF-Q53G59-F1-v4), PATZ1 (pdb: 6GUV) monomer and dimer. Point mutations for KLHL12 (L20D/M23D/L26D/L49K) are marked in red within the KLHL12 AlphaFold prediction. Overlay and alpha carbon-root mean square deviation (C_α_ RMSD) show structural similarity between proteins. The fold is especially conserved for the dimerization interface which has a C_α_ RMSD of 0Å (dark blue) between different proteins. Differences larger than 3Å are observed for N- and C-termini and for the loop region between the beta-strand B3 and α-helix A2. In the case of PATZ1 the B3 β-strand couldn’t be structurally resolved due to an additional large flexible linker between the α-helix A2 and β-strand B3, which adds to the poorer C_α_ RMSD within that region for the comparison between PATZ1 and KLHL12 and between PATZ1 and KEAP1.

**Supplementary Figure 2.**
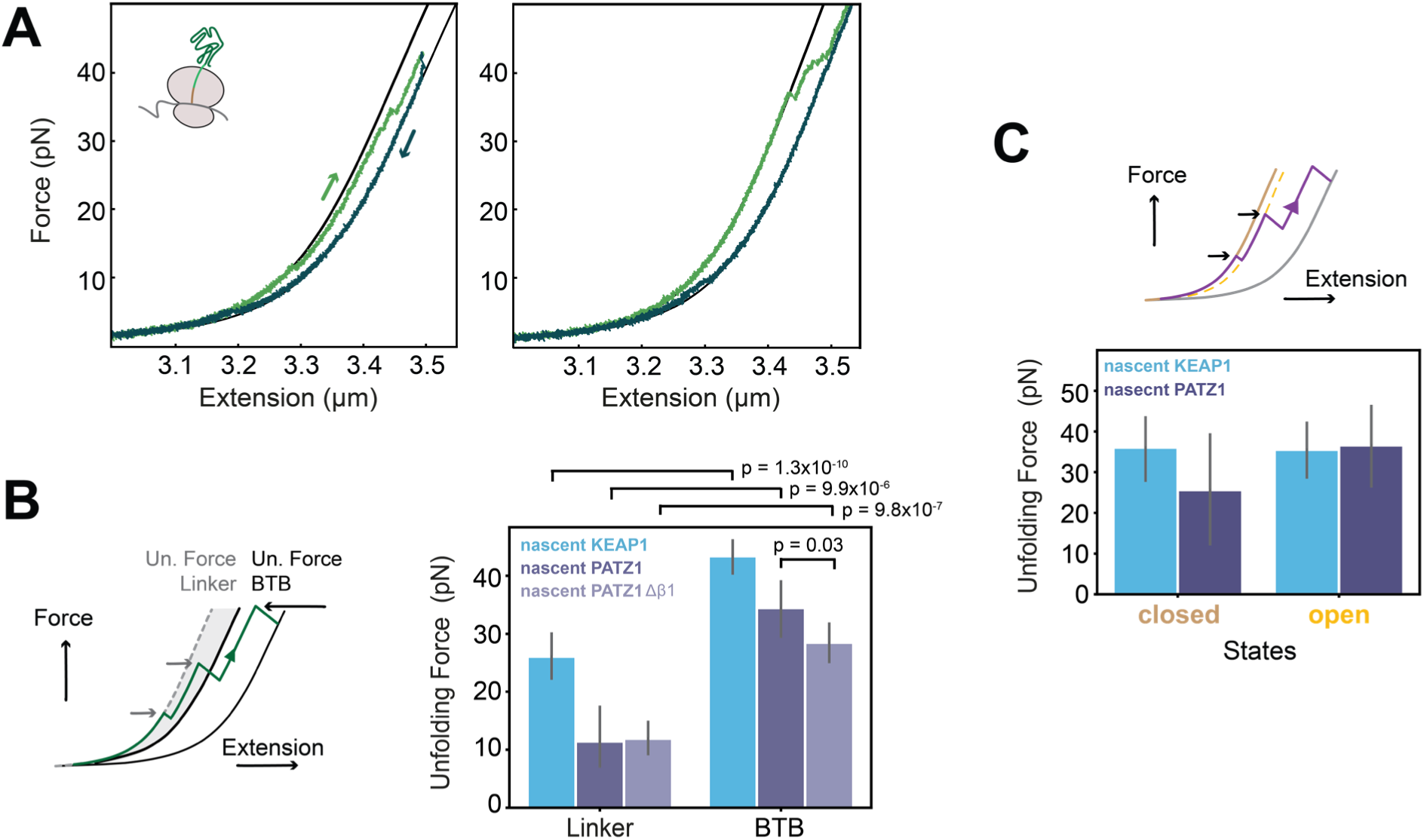
A) Example force-extension traces of a ribosomal nascent chain construct (50-196 aa) of nascent KEAP1 without artificial linkers showing full compaction. Owing to 17 amino acids of the natural sequence at the C-terminus of the BTB domain in addition to the SecM strong sequence, the full BTB domain is exposed outside of the ribosomal tunnel to allow for complete folding. Both traces show how the BTB monomer is initially compacted (left worm-like chain model) during pulling and unfolds stepwise to the calculated length of the fully unfolded length of the monomer (right worm-like chain model). B) Unfolding forces for states at contour lengths corresponding to linkers and BTB domain defined. Since the here artificial linkers may not form stable folds and only weak interactions, those interactions would break and unfold first. Thus, we considered the unfolding forces of the contour lengths corresponding to the linker (grey area in cartoon on the left) and compared them with the unfolding forces of states corresponding to BTB unfolding. Indeed, a significantly lower mean unfolding force (p < 0.05) was observed. Error bars: 95% confidence interval. See methods for N-values. C) Measured unfolding forces to partially unfold the closed and open BTB monomer states. We used a +/− 2.5 nm margin around the lengths of these states and quantified the unfolding forces (black arrows in the cartoon). Mean unfolding forces were quite similar for both states and relatively high (closed nascent PATZ1: 25.6 pN, closed nascent KEAP1: 36 pN, open nascent PATZ1: 36.6 pN, open nascent KEAP1: 35.5 pN), indicating a stability of the fold. Error bars: 95% confidence interval. See methods for N-values.

**Supplementary Figure 3:**
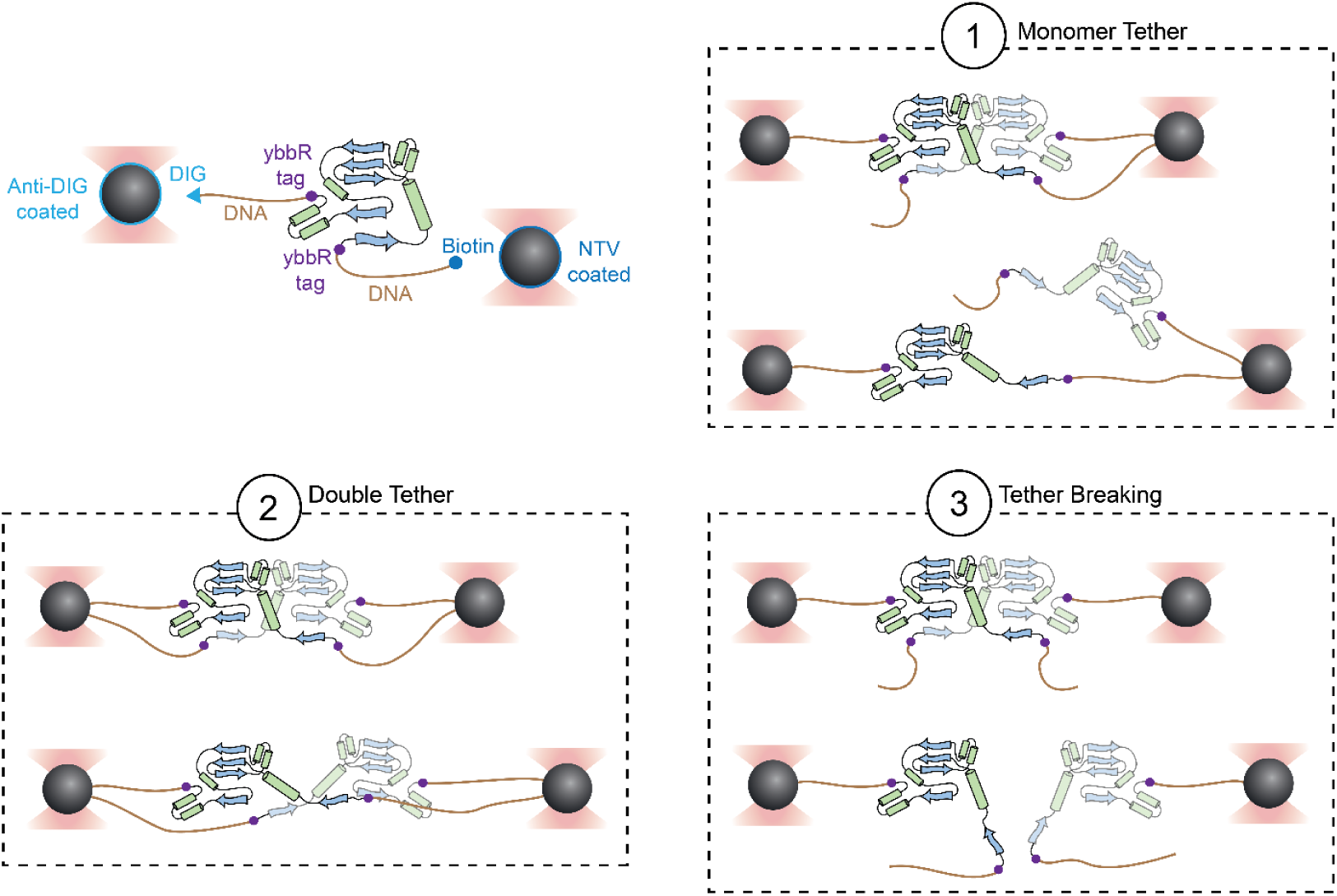
Purified BTB monomer tethering. Visualization of the different tethers that can form and how we obtain tethered monomers (nr. 1, top-right). YbbR tags were genetically introduced at each terminus of the BTB monomer, coupled to 20 nucleotide-long oligos, modified with coenzyme A using Sfp synthase (Sfp 4’-phosphopantetheinyl transferase), and then covalently ligated to DNA tethers. Tethering naturally selects only those constructs where one DNA handle has a biotin and the other has a digoxigenin, as constructs with two of the same handles cannot form a tether between the two beads. This DNA attachment allows one to select for BTB monomers (nr. 1) in our optical tweezer assay, even though up to 4 DNA handles can in principle attach to the 4 termini of purified dimers. Specifically, as visualized in the cartoons, the first stretching curve will either result in: 1) a single BTB monomer tethered at its termini between the beads, 2) two BTB monomers each tethered individually between the two beads, or 3) tether breakage. Case 3 is readily recognized by a sudden complete rupture event in force-extension curves during the first pull that breaks the tether. In case 2 a double tether is formed. It can be identified by the absence of an overstretching plateau at 65pN, owing to the fact that two tethers share the mechanical load and hence can resist a higher load. Only a single monomer tether (case 1) shows an overstretching plateau at 65 pN due to DNA melting.

**Supplementary Figure 4:**
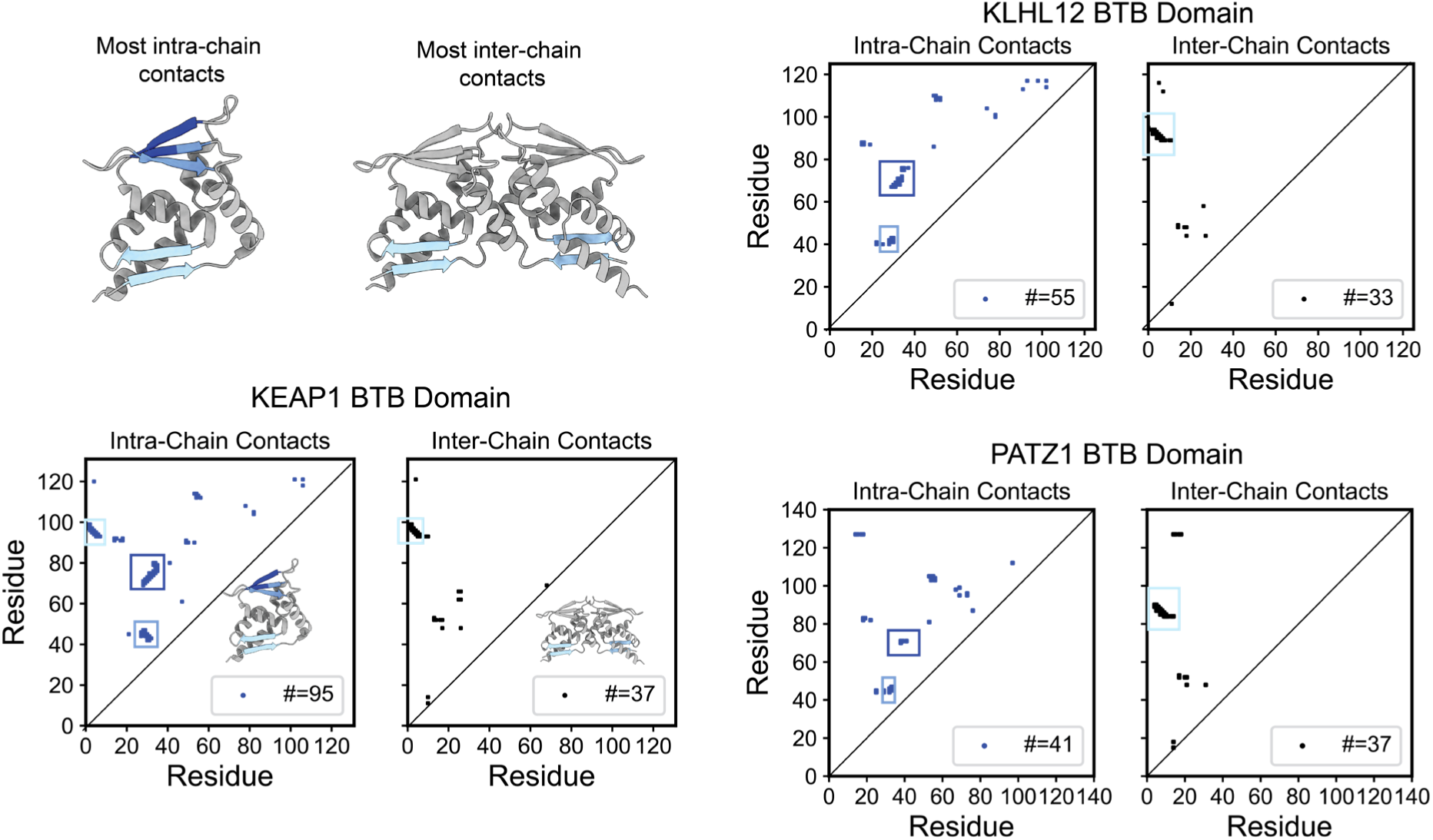
Intra- and inter-chain contacts for BTB domains. Intra- and inter-chain contacts of the studied BTB monomers and dimers. Two residues were considered in contact if their corresponding carbon alpha atoms were in a distance smaller or equal to 7Å from each other. Atom positions are taken from the same crystal structures as indicated in **Fig. S1A**, apart from the closed monomer structure of KEAP1 (pdb: 6w66). Many contacts are found between beta sheets, which are marked and highlighted correspondingly in blue within the structure.

**Supplementary Figure 5:**
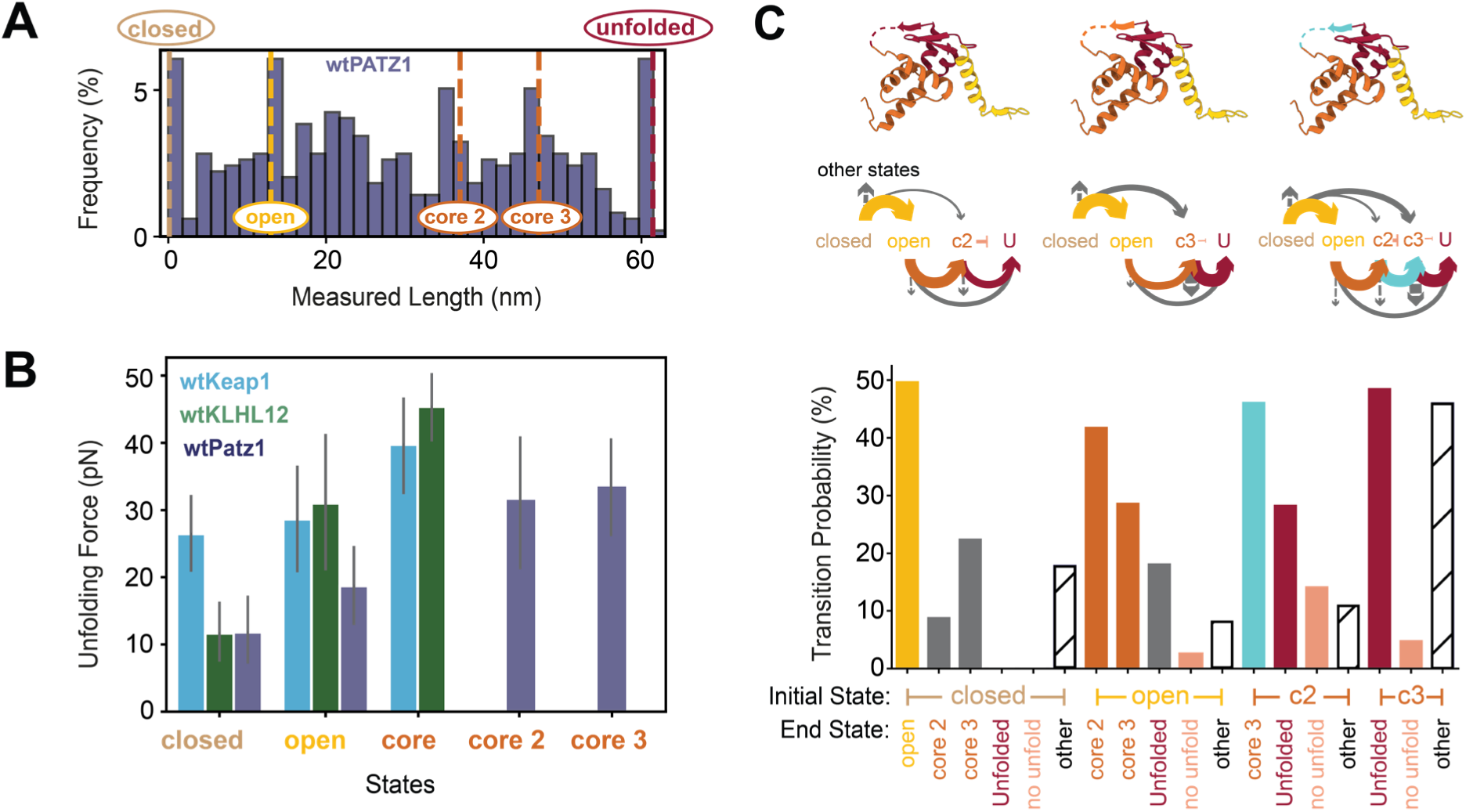
Extended analysis of purified PATZ1 monomer. A) Contour length histogram of purified wtPATZ1 monomer (as in Fig. 2C) with dotted lines indicating the most pronounced contour length states (partial folds). Due to PATZ1’s unique A2/B3 loop, it does not map onto the “core” state as seen for wtKEAP1 and wtKLHL12. Rather, two alternative “core” states match the marked peaks in the histogram: one in which the flexible loop unfolds with the c-terminal region (“core 3”) and one where only the c-terminal region unfolds but the loop not (“core 2”). See methods for N-values. B) Mean unfolding forces for wtPATZ1 “core 2” and “core 3” states added to the unfolding forces of wtKEAP1 and wtKLHL12 states as displayed in Fig.2E. wtPATZ1 core states show lower unfolding forces in contrast to the unfolding forces of wtKEAP1 and wtKLHL12 core states (p = 0.01), which is consistent with decreased stability due to fewer contacts within that region (see **Fig. S4**). Error bars: 95% confidence interval. See methods for N-values. C) Transition probabilities (see methods) between defined states for wtPATZ1 monomer. The colored structures at top indicate the possible unfolding sequences. (1) left: the flexible loop is part of the partial fold which unfolds last (red), giving rise to the intermediate “core 2” (2) middle: the flexible loop unfolds with the rest of the C-Terminus (orange), giving rise to the intermediate “core 3”. (3) the unfolding trajectory contains both intermediates, “core 2” and “core 3” with an additionally unfolding step between “core 2” and “core 3” (flexible loop – cyan). See methods for N-values.

**Supplementary Figure 6:**
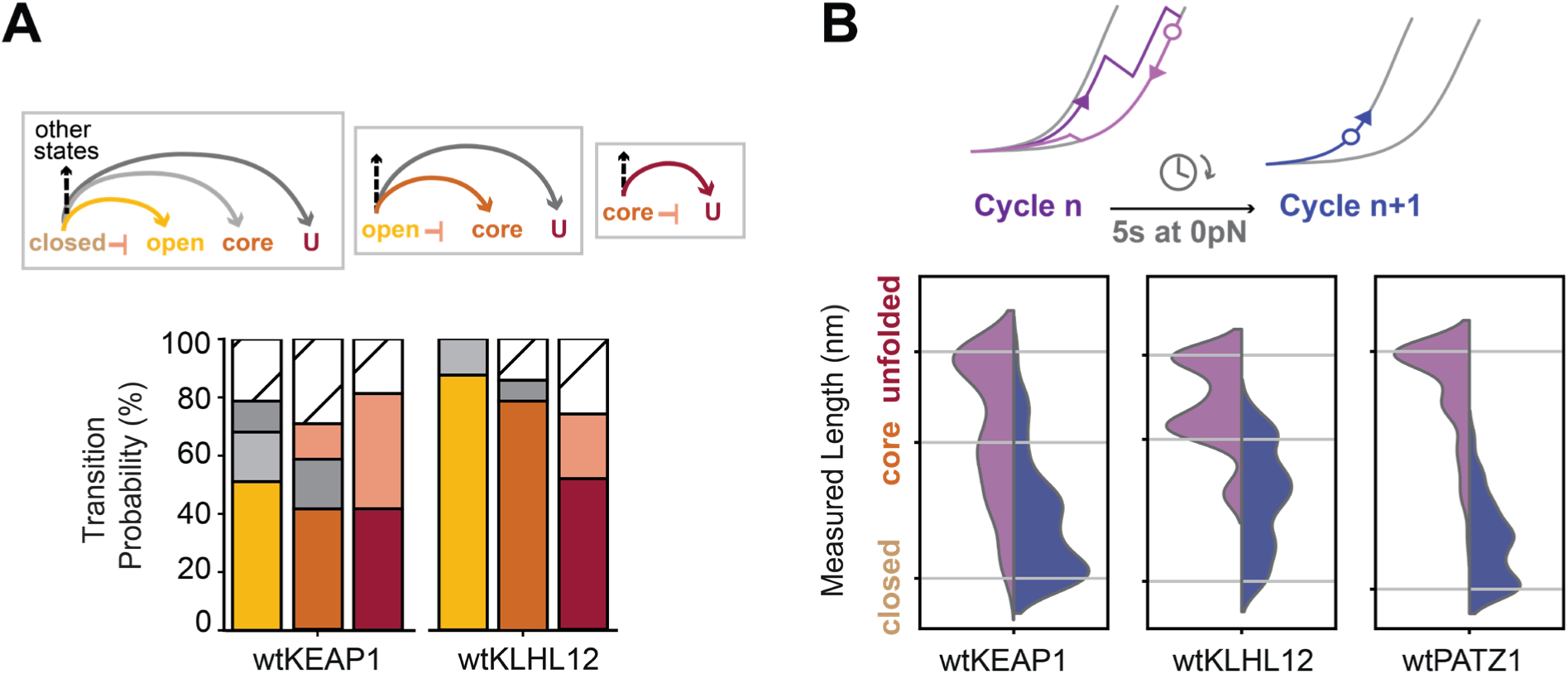
Unfolding and refolding of purified BTB monomers. A) Transition probabilities between states, as measured using optical tweezer experiments on purified BTB monomers (see main Fig. 2). For molecules in the closed, open or core states (lengths in a +/− 3nm window around their expected lengths, see methods), we scored transitions to one of these states of larger length during stretching, or any other state of different length, or remained in the same state without length changes (see cartoons, top, and **Fig. S5C**). Stacked bars indicate transition probability or frequency corresponding to the cartoon arrows. For each protein, first cartoon corresponds to first stacked bar, second cartoon to the stacked bar in the middle and the last cartoon to the third stacked bar. Both proteins showed a similar most prevalent unfolding trajectory, which follows a consecutive sequence of the defined states from closed to open to core and then either remained in the core state or unfolded to the fully unfolded state. See methods for N-values. B) Refolding was studied by quantifying the most unfolded state within one stretch-relax cycle (cartoon: purple circle) and the state populated after a waiting period at 0 pN (cartoon: blue circle) to allow for compaction without a counteracting force. The most unfolded states (purple) and the most compacted states (blue) were plotted on a vertical axis with a kernel density estimation curve. For wtKEAP1 (216 events) and wtKLHL12 (64 events), the most unfolded states within one stretch-relax cycle were often either the fully unfolded state or the “core” state, consistent with our transition probability analysis of panel A). The most unfolded state for wtPATZ1 (128 events) was most often the fully unfolded state, which is in line with the decreased stability of the core states (see also **Fig. S5 & S4**).

**Supplementary Figure 7:**
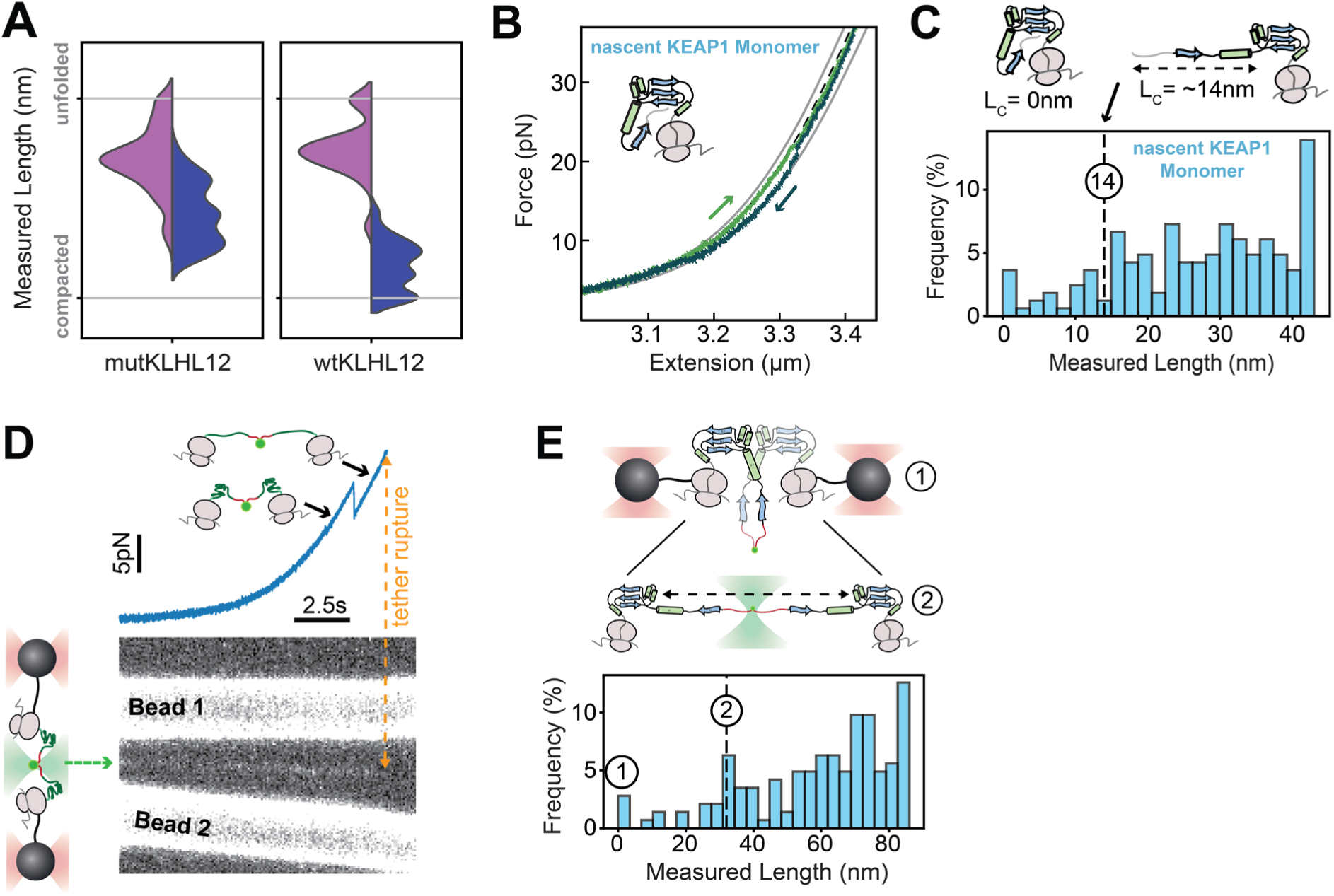
BTB monomer and dimer characterization. A) Refolding analysis as described in Fig. 3C and **Fig. S6B**, here for the wtKLHL12 fusion dimer (52 events) and its dimerization defective mutant. A similar distribution for the most unfolded states per cycle (purple) for both proteins is observed, indicative for similar partial folds for both proteins. Consistently, in contrast to the dimerization defective mutant, the fully compacted state is only observed for the wild-type version of the wtKLHL12 BTB domain, indicating dimer formation. B) Example Force-Extension trace of a stretch-relax cycle for the BTB monomer in an incomplete translation stage (50-159 aa), which misses the C-terminus inclusive the beta-strand B4 that β1 folds back onto. However, all structural elements of the core partial fold are translated. C) Contour length histogram of BTB monomer in incomplete translation stage (50-159aa) as in B (165 states). Compacted states are observed with a low frequency, in line with α1 and β1 and the N-Terminal flexible linker being unfolded (see cartoon). Partial folds bigger than 14 nm, corresponding to the core state being (partially) folded, are more frequently observed. D) To prevent tether loss nascent chains are linked up by a FlAsH dye^3^, which binds to a bipartite tetracysteine motif formed by two engineered cysteines at the end of a linker at each nascent chain N-terminus. Cartoon left: Bound FlAsH becomes fluorescent and is detected by scanning a confocal excitation beam (green) along the molecular tether. Corresponding data (right) of detected fluorescence scans in time (bottom) and measured force-time trace (top). The signal between beads corresponds to FlAsH fluorescence. Dashed line indicates tether rupture, which correlates with the loss of fluorescent signal. E) Contour length histogram of the incomplete translation RNC construct pairing (as panels B and C) (143 states). Dotted lines and numbers indicate contour length states matching the cartoon at top.

**Supplementary Figure 8:**
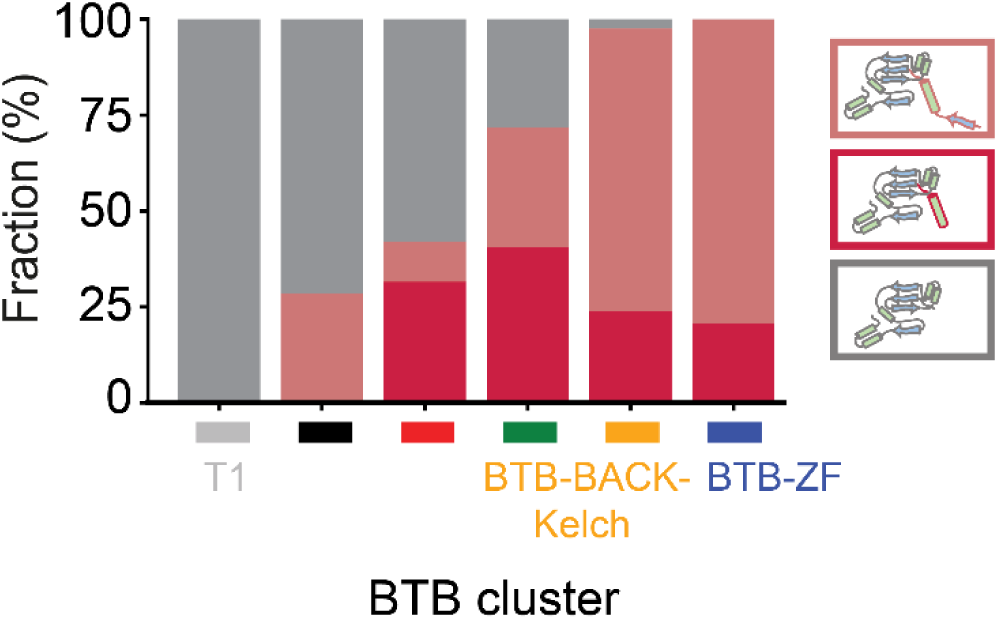
Analysis of structural features across BTB proteome. Structural analysis of the amino-terminal extension for BTB proteins per cluster (see Fig. 4A). Alpha-fold^2^ was used to predict structures and to assess the amino-terminal extension. Only structures with high confidence of the BTB domain region were used. Proteins were categorized in having either no amino-terminal extension (grey), an amino-terminal extension of α1 (dark red) or an amino-terminal extension of both α1 and β1 (light red). Superfamily clusters of the BTB-ZF, BTB-BACK-Kelch predominantly consist of BTB proteins containing an amino-terminal extension.

